# Distinct clonal evolution of B-cells in HIV controllers with neutralizing antibody breadth

**DOI:** 10.1101/2020.09.02.277566

**Authors:** Deniz Cizmeci, Giuseppe Lofano, Anne-Sophie Dugast, Dongkyoon Kim, Guy Cavet, Ngan Nguyen, Yann Chong Tan, Michael S. Seaman, Galit Alter, Boris Julg

**Affiliations:** Ragon Institute of MGH, MIT and Harvard, Cambridge, MA 02139, USA; Atreca Inc, Redwood City, CA, USA; Esco Ventures X, Singapore 139951; Center for Virology and Vaccine Research, Beth Israel Deaconess Medical Center, Boston, MA 02115, USA

**Author notes:** corresponding author: Boris Julg, MD PhD, The Ragon Institute of MGH, MIT, and Harvard, 400 Technology Sq, Cambridge, MA 02139.

**Keywords:** neutralizing antibodies, HIV controller, V(D)J rearrangement, clonal selection

## Abstract

A minor subset of individuals infected with HIV-1 develop antibody neutralization breadth during the natural course of the infection, often linked to chronic, high level viremia. Despite significant efforts, vaccination strategies have been unable to induce similar neutralization breadth and the mechanisms underlying neutralizing antibody induction remain largely elusive. Broadly neutralizing antibody responses can also be found in individuals who control HIV to low and even undetectable plasma levels in the absence of antiretroviral therapy, suggesting that high antigen exposure is not a strict requirement for neutralization breadth. We therefore performed an analysis of paired heavy and light chain B-cell receptor repertoires in 12,591 HIV-1 Envelope-specific single memory B-cells to determine alterations in the BCR immunoglobulin gene repertoire and B-cell clonal expansions that associate with neutralizing antibody breadth in 22 HIV controllers. We found that the frequency of genomic mutations in IGHV and IGLV was directly correlated with serum neutralization breadth. The repertoire of the most mutated antibodies was dominated by a small number of large clones with evolutionary signatures suggesting that these clones had reached peak affinity maturation. These data demonstrate that even in the setting of low plasma HIV antigenemia, similar to what a vaccine can potentially achieve, BCR selection for extended somatic hypermutation and clonal evolution can occur in some individuals suggesting that host-specific factors might be involved that could be targeted with future vaccine strategies.

## Introduction

The induction of cross-reactive broadly neutralizing antibodies (bNAbs) that can effectively neutralize a majority of circulating HIV-1 strains is a major goal of current HIV vaccine development, but no vaccine candidate to date has achieved this goal (1). During natural HIV-1 infection, development of neutralizing antibody responses is seen in about 10-30 % of HIV-1-infected individuals but often requires years of infection (2). Specifically, high viremia (3, 4) with rapid viral diversification (5, 6) along with infection-associated immune activation (7) have been associated with the evolution of antibody neutralization breadth. One specific feature that characterizes many bNAbs is the high mutation frequency in their variable domains, indicative of many cycles of somatic hypermutation (SHM) in Ig variable-diversity-joining (V(D)J) gene segments, ultimately resulting in superior B cell receptor-antigen affinity maturation (8). The requirement for repeated cycling through the germinal centers (GC) to accumulate affinity and specificity enhancing mutations is a lengthy process and is reflected by the time required for bNAbs to be elicited in the presence of a constantly evolving virus. However, bNAbs with high potency and exquisite breadth against cross-clade viral isolates have also been isolated from HIV-infected individuals who spontaneously control HIV to low plasma levels in the absence of antiretroviral therapy (“HIV controllers”) (9, 10). In these individuals, evolution of neutralization breadth was linked to low but persistent HIV viral antigenemia in the setting of a unique inflammatory profile, while antigenic diversity was not a significant contributor (11). Thus, neutralizing antibody breadth may be achievable in the absence of high levels of viremia and rapidly evolving viral diversity, therefore suggesting a scenario that more likely can be accomplished with vaccination strategies. In this study, we selected 22 controllers who exhibited different levels of neutralization breadth (ranging from 0/11 to 11/11 of tier 2/3 viruses neutralized) (11) to eludicate features of the HIV-specific antibody repertoire that are associated with neutralization breadth. We performed a natively-paired heavy and light chain B cell receptor repertoire analysis of HIV-1 Env-specific memory B-cells using high-throughput single-cell sequencing to determine alterations in the BCR immunoglobulin gene repertoire and B-cell clonal expansions that track with neutralization breadth.

## Results

### Study cohort characteristics

22 HIV-infected individuals, 4 female and 18 male, were included in this study. 11 participants were African American and 11 were of European descent. All individuals were US based and therefore likely infected with clade B viruses. Plasma samples were tested for neutralizing activity using a standard reference panel of tier 2 and 3 clade B Env pseudoviruses (12). Six individuals neutralized >90% of viruses at 50% inhibitory dose (ID50) titers above background, and were classified as Top-Neutralizer (TN), while 7 individuals neutralized <10% of the panel viruses and were classified as Non-Neutralizer (NN). The remaining individuals were considered intermediate-neutralizer, exhibiting neutralization of 36% to 82% of the virus panel (Supplemental Table 1). There was no statistically significant difference in age (mean 53 years; min 44, max 63), CD4 count (mean 726 cells/μl (SD 168), HIV RNA levels (mean 961 copies/mL (SD 1645) and time-off-ART (mean 17 years; min 4.5, max 30) between the TN and NN groups (Supplemental figure 1). Single HIV-1 Env-specific CD19^+^CD20^+^IgM^-^IgA^-^ memory B-cells (MBC) were FACS-sorted (Supplemental figure 2) and we obtained repertoires of natively-paired, full variable region IGH and IGL by Immune Repertoire Capture® analysis. 12,591 total sequence pairs were generated for all study participants, with 5,771 sequences from TN, and 2,707 sequences from NN. To exclude potential bias caused by the number of input cells or sequences, we normalized the proportion of each repertoire signature across all sequences in each individual (see methods). TN memory B-cells showed an Ig subclass distribution similar to serum IgG antibodies, with average levels of IgG1 (78.6%) > IgG2 (12.5%) > IgG3 (8.8%) > IgG4 (0.1%) while Ig subclasses from NNs differed in hierarchy with average levels of IgG1 (82.5%) > IgG3 (11.1%) > IgG2 (6.2%) > IgG4 (0.2%) (Fig. 1A). As IgG3 is often one of the earliest subclasses to be elicited during infection (reviewed in (13)) being the first IGHG gene located within the locus (14), the higher percentage of IgG3 in NN versus TN might suggest a higher proportion of more recently induced MBCs in the NNs.

**Figure 1:**
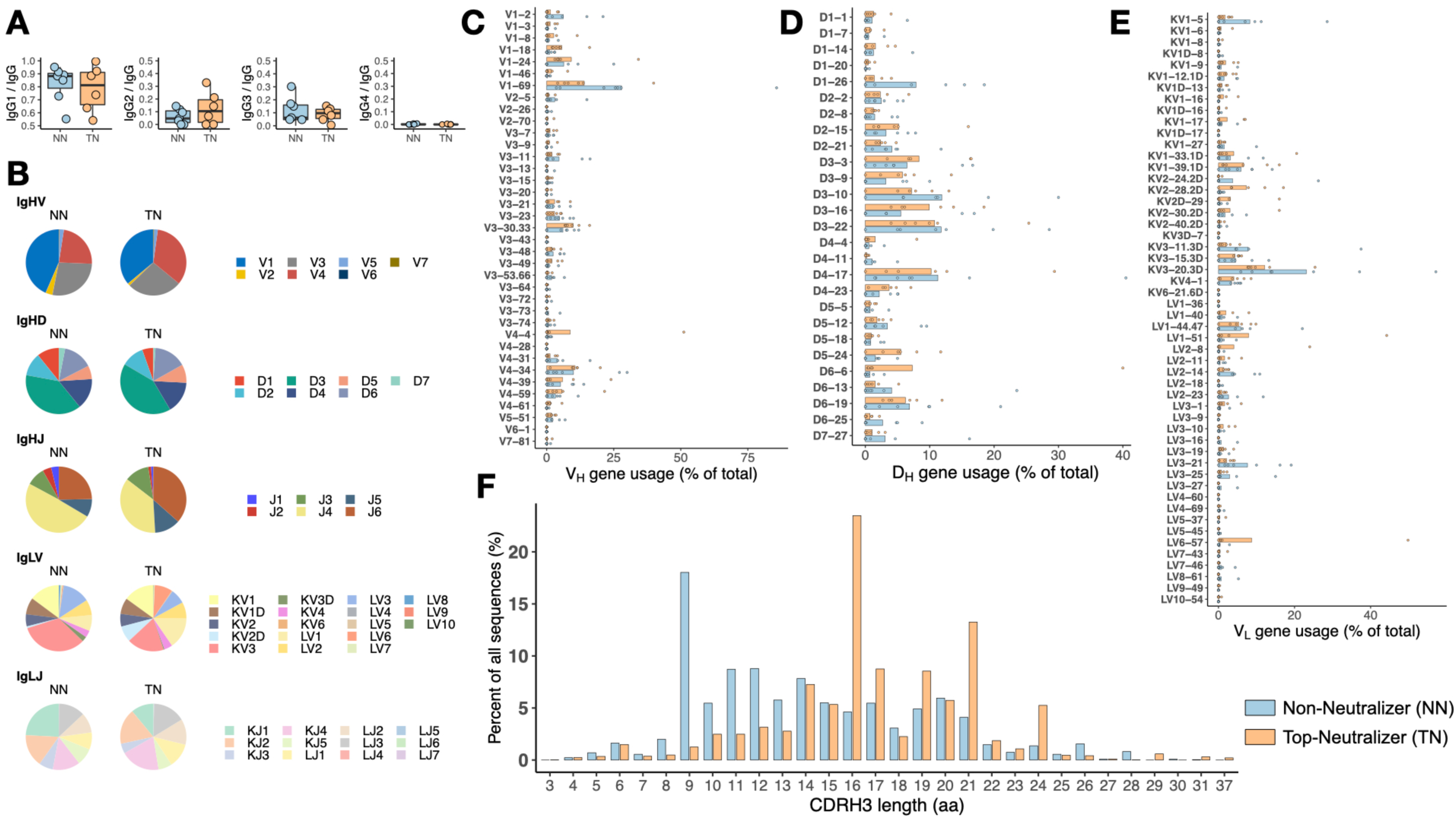
Non-Neutralizer and Top-Neutralizer MBCs with similar BCR repertoires. (A) IgG subclass distribution in NNs and TNs. (B) The frequencies at which the IGH V/D/J and IGL V/J gene families are expressed by Non-Neutralizer (NN) and Top-Neutralizer (TN) MBCs are shown in pie charts as the average of percentages of gene sequences obtained in each individual. The mean frequency of individual VH (C), DH (D) and VL (E) gene segments in NN and TN MBCs are shown as a percentage of all annotated genes. Shown are means. (F) CDRH3 length distribution (in aa) in all NN and TN sequences pooled together.

### IGH-VDJ and IGL-VJ Gene Segment Usage

The MBC repertoires were found to use 40 IGHV genes representing all 7 human VH gene families. Of the seven IGHV gene families, IGHV1 was most commonly used in both NN and TN repertoires, consistent with prior reports that anti-HIV antibodies frequently use IGHV1 genes (15, 16). The top three most frequently used IGHV genes were IGHV1-69, IGHV4, and IGHV1-2 in both the TN and NN repertoire, followed by IGHV4-4 in the TN and IGHV1-2 in the NN repertoire (Fig. 1C). Interestingly, IGHV1-69, which is enriched in neutralizing antibodies that target HIV-1 V2 and gp41 as well as other viral infections such as influenza and Hepatitis C virus (17-19), was significantly less frequently used in TN than in NN (mean percentage 14% vs 28%, *p* < 0.001 two-way ANOVA followed by Sidak’s multiple comparisons test). Furthermore, IGHV1-2, which is utilized by CD4bs antibodies such as VRC01, 3BNC117 and N6 (20-22) as well as in V3-glycan dependent bNAbs (23) and gp120-gp41 interface bNAbs, was more frequently found in the NN group than in the TN group, suggesting that the selection of this VH gene by itself did not predict a higher likelihood for bNAb development. For IGHD and IGHJ genes no statistical significant differences were observed. The most frequently used heavy chain VDJ combinations were IGHV1-69, IGHD3-22, IGHJ4 in NN and IGHV1-24, IGHD4-17, IGHJ4 in TN. Among IGLV and IGLJ genes, IGLKV3 and IGLKV3-20 dominated in both TN and NN repertoire (Fig. 1E). This is consistent with previously reported data showing the dominance of IGLKV3-20 (24). IGLKV3-20, although preferred in both TN and NN repertoires, was used significantly more frequently in the NN compared to the TN repertoires (mean percentage 21% vs 12%, *p* < 0.01 two-way ANOVA followed by Sidak’s multiple comparisons test). Taken together, these results suggest that the VDJ gene usage was similar overall in the IGH repertoires of TN and NN, and the frequencies of germline genes that have been associated with known bNAbs did not associate with neutralization breadth in our cohort.

### Characteristics of CDRs

CDRs play critical roles in the binding of antibodies to antigens and the length of CDR3 has been a characteristic feature of certain bNAbs (25). The CDRH3/L3 length in TNs was between 3 and 37 aa and 5 and 17 aa, respectively and 4 to 31 aa and 5 to 14 aa in the NNs, respectively. While the median CDRH3 length was slightly longer in the TN repertoires than the NN repertoires (16 vs 14 aa, respectively) this difference did not reach statistical significance (*p* > 0.05 Wilcoxon rank sum test; Fig. 1F). Furthermore, no significant differences in length were observed between the groups for either CDRH1/L1 and CDRH2/L2.. The CDRH3 lengths observed here are more consistent with what has been described in the CD4 binding site specific bNAbs like VRC01 and 3BNC117, which have CDRH3 lengths of 14 and 12 respectively, while V2 and V3 glycan specific bNAbs like PGDM1400 and PGT121 have substantially longer CDRH3 length with 34 and 26 aa, respectively (25).

### Rates of somatic hypermutation in IGHV and IGLV

When we plotted the number of somatic hypermutations in IGHV (Fig. 2A), we noticed that the TN repertoires had significantly higher frequencies of nucleotide mutations compared to NN (Mean ± SD: 54 ± 17 mutations for TN and 29 ± 8 mutations for NN, respectively, *p* < 0.05 unpaired t-test) with more than 65% of sequences showing mutation counts greater than 60. When comparing the frequency of mutations in IGHV (TN, NN and intermediate neutralizers) with the degree of serum neutralization breadth across the entire cohort, a direct correlation was observed (Spearman’s rho = 0.60, *p* < 0.01) (Fig. 2B and C). Similarly, somatic hypermutation in IGLV (Fig. 2D) was more frequent in TN than NN (Mean ± SD: 35 ± 18 mutations for TN and 14 ± 5 mutations for NN, respectively, unpaired t-test *p* < 0.05) with more than 50% of sequences showing mutation rates greater than 40. Here as well, the frequencies of mutations in IGLV correlated with neutralization breadth (Spearman’s rho = 0.63, *p* < 0.01; Fig. 2E and F). We were next interested to determine whether the IGH and IGL V-J gene usage would differ between the more (with more than 60 mutations in IGH, 40 mutations in IGL) and less (less than or equal to 60 mutations in IGH, 40 mutations in IGL) mutated sequences. While including all sequences, independent of the mutation rates, there was no apparent difference in IGHV-J (Fig. 2G) or the IGLV-J combinations between NNs and TNs (Fig. 2H). For the sequences with more than 60 mutations in IGHV, however, which occurred nearly exclusively in the TNs, the pattern was distinct, demonstrating a selection for fewer dominant V-J combinations (Fig. 2G). In particular JH4/VH1-24, JH6/VH4-4, JH6/VH3-30.33, and JH6/VH4-34 were significantly enriched in TN sequences with high mutation rates (*p* < 0.01, one sided Fisher exact test followed by correction for multiple comparison using Benjamini-Hochberg method). The enrichment of HV4-34 usage (*p* < 0.001, one sided Fisher exact test followed by correction for multiple comparison using Benjamini-Hochberg method), a feature of autoimmunity (26), in the TN sequences with high mutation rates, is consistent with a recent report that suggested that HIV infection induces a permissive state in which potentially autoreactive clones that would otherwise be eliminated can persist (27). A similar contraction to fewer dominant V-J combinations was also observed for sequences with more than 40 mutations in IGLV (Fig. 2H). Assuming that the somatic IGHV mutation rates in B cells correlate with the total exposure time in the GC environment, and in light of the fact that the TN and NN did not differ in age, time-off-ART and viral loads, our data suggests that in the TNs, B-cell clones were selected for extensive SHM.

**Figure 2:**
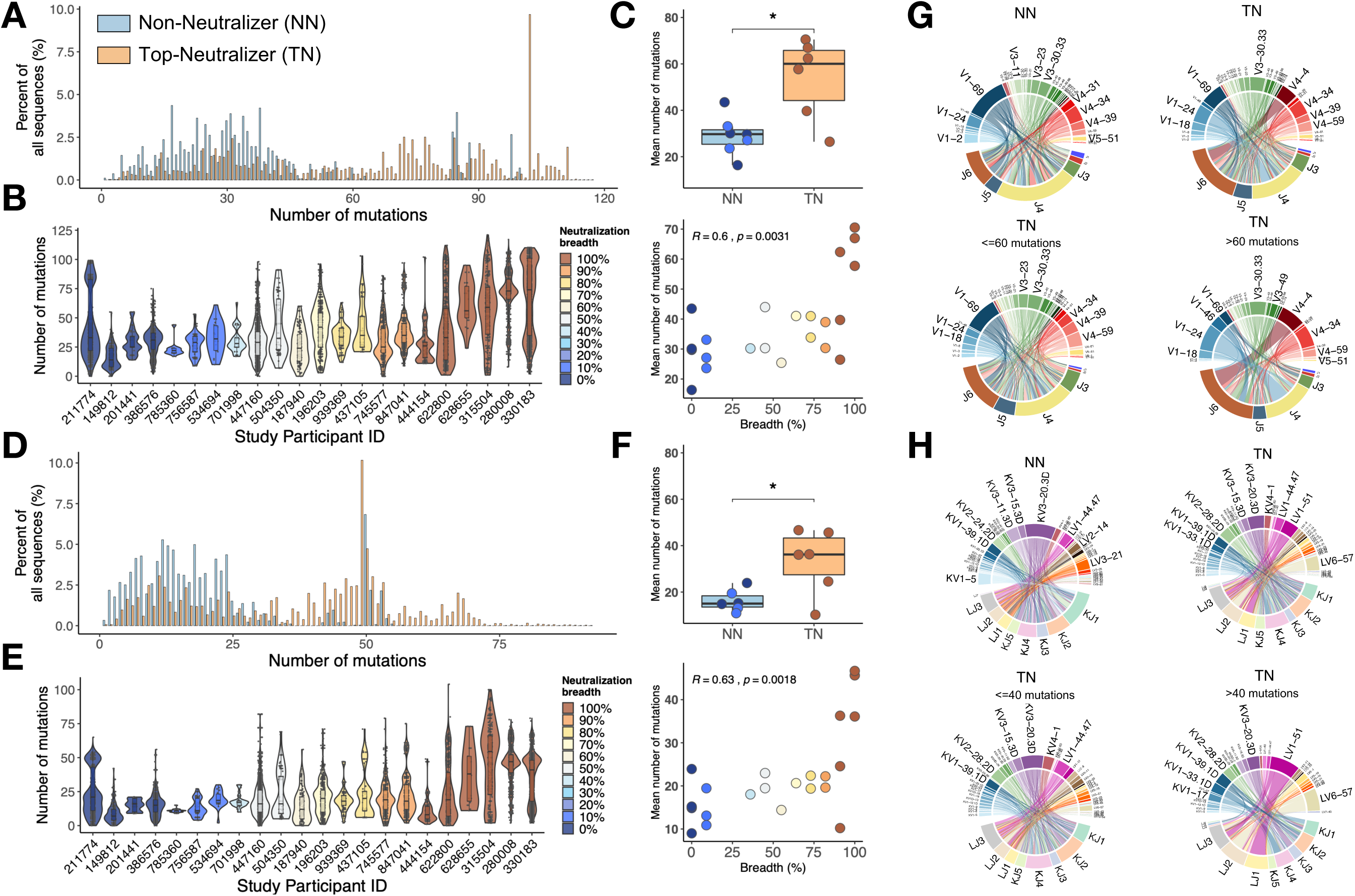
Mutation characteristics of IgHV and IgLV genes and associations with neutralization breadth. **(A)** The overall mutation frequencies in IGHV in TNs versus NNs are significantly different with mean number of mutations of 54 in TNs and 29 in NNs (unpaired t-test *p* < 0.05). **(B)** The number of mutations in IGHV for each of the 22 study participants is plotted and the serum neutralization breadth is color coded ranging from blue (0% neutralization) to red (100% neutralization). **(C)** The mean frequency of mutations in IGHV correlates with the serum neutralization breadth in % (Spearman’s rho = 0.60, *p* < 0.01). Similarily, for IGLV, the mean number of mutations differ (35 in TNs and 14 in NNs **(D)** and correlates with the serum neutralization breadth in % (r = 0.63, *p* < 0.01) **(E and F)**. Circos plots summarize the combinations of V and J segments used in the rearranged IGH genes **(G)** and IGL genes **(H)** expressed by MBCs. Top circos graphs demonstrates V-J combinations expressed by NN MBCs and TN MBCs independent of the number of detected mutations. Bottom circos graphs demonstrate V-J combinations expressed by TN MBCs with low and high mutation rates. For each plot, the bottom half depicts J genes and the top half depicts Vgenes. To exclude potential bias caused by the number of input cells, the number of sequences in each repertoire signature was weighted by the total number of sequences in each individual (see methods). The arc length of each segment denotes the normalized frequency at which each gene segment was identified. Rearrangement of a J gene with a V gene segment in a clonal Ig sequence is represented by a ribbon (ribbons carry the color of the HV or LV family of the gene participating in the pairing). The width of the ribbons corresponds to the weighted frequency at which each particular HV-HJ or LV-LJ rearrangement was used in the respective MBC repertoire.

### Clonal selection

To further assess the clonal nature of the BCR repertoire in TNs, we selected subjects (subject IDs: 330183, 280008, 622800) for whom a total of more than 600 sequences were available. Sequences were clustered into clones using single-linkage clustering. Each unique clone is a cluster of sequences that meet the following criteria (1) derived from the same individual; (2) share the same V and J gene segment annotations in heavy chain and light chain (3) have equal CDRH3 length; and (4) CDRH3 aa sequences match with =/> 90% similarity. This clonal assignment identified 703 clones (mean number of sequences 8.4) and showed that larger sized clusters (dominant clones) consist of highly mutated sequences (Fig. 3A). To quantify the greater clonality of the highly mutated sequences, we used an unevenness measure, the Gini index. Sequences were divided into two sets, highly mutated (>60 mutations) and less mutated (=<60 mutations) sequences. Each set of sequences was clustered into clones and the Gini index was calculated based on the clone size distributions. For all three subjects, the Gini index was greater in the highly mutated set (*p* < 0.05 paired t-test), thus supporting the observation that the BCR repertoire becomes more narrow with increasing mutation rates, suggesting the dominance of certain clones (Fig. 3B).

**Figure 3:**
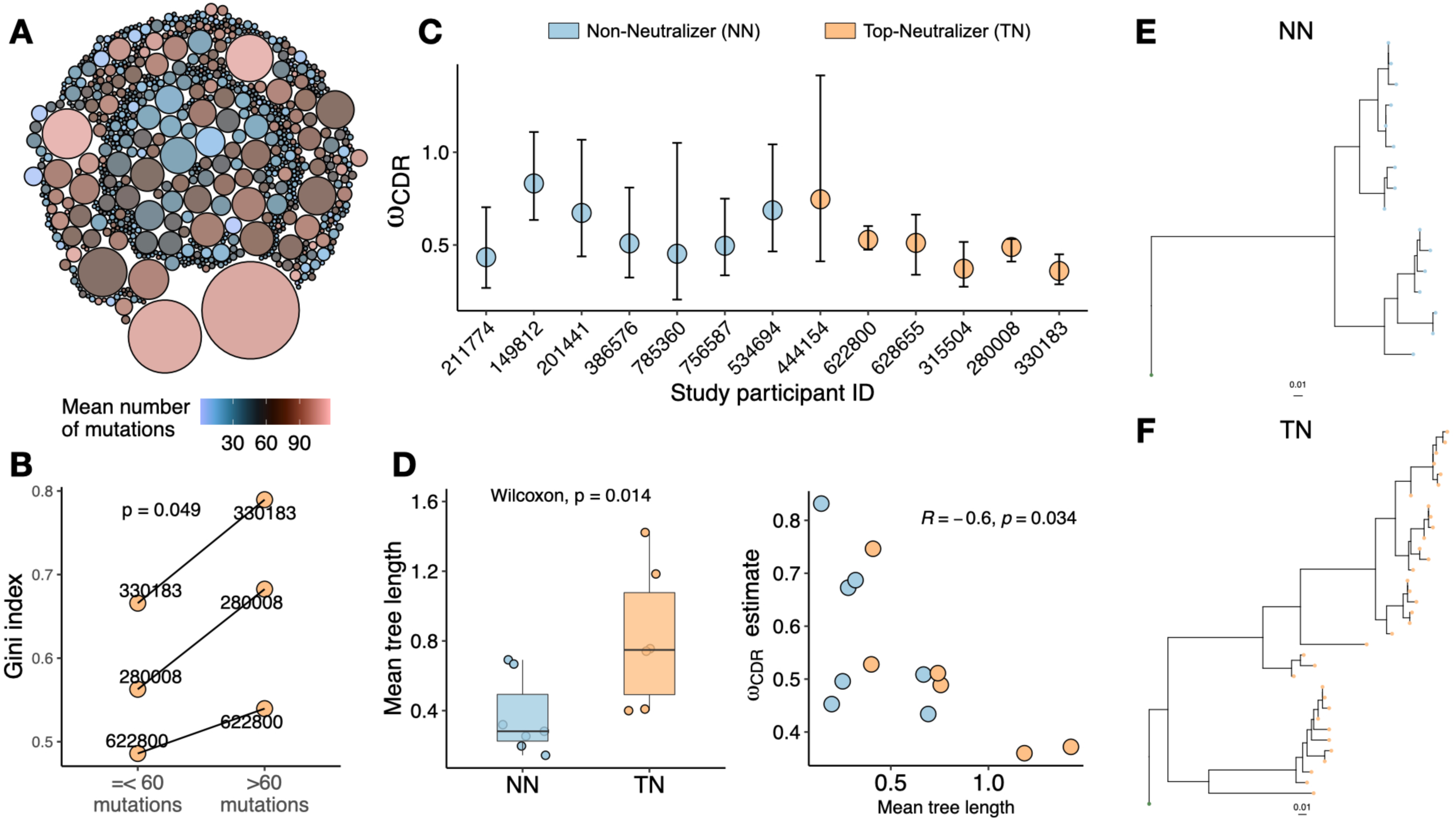
Clonality analysis. A total number of 703 clones were obtained. **(A)** Each clone is represented as a circle. The sizes of the circles are scaled to the number of sequences in each clone (mean number of sequences 7.7); the colors represent mean number of mutations per clone. Dominant clones consist of highly mutated sequences. **(B)** Gini index (0 represents perfect equality and 1 perfect inequality) was calculated as an estimate of clonality for TNs for which a total of more than 600 sequences were obtained (subject IDs: 330183, 280008, 622800). Per each subject, sequences were divided into two sets as high mutated sequences (>60 mutations) and low mutated sequences (=<60 mutations). Each set of sequences were clustered into clones and Gini index was calculated. Gini Index is significantly higher (paired t-test, *p* < 0.05) in highly mutated sequence sets supporting the presence of fewer dominant clones. **(C)** Maximum likelihood point estimates of ω with 95% confidence intervals. **(D)** Estimates of mean tree length (total substitutions per codon within a lineage, averaged across all lineages within a repertoire) were compared between TN and NN using Wilcoxon rank sum test (left graph). The correlation between ω and mean tree length calculated using Spearman correlation. Largest lineage trees in NNs **(E)** and TNs **(F)**.

Next, we estimated the evolutionary process of mutation and selection in B cells using phylogenetic models. Selective dynamics have been estimated using ω: dN/dS, the ratio of nonsynonymous (amino acid replacement) and synonymous (silent) mutation rates (28, 29). Low ω_CDR_ values indicate fewer amino acid changes in CDRs than expected suggesting negative BCR selection to remove affinity-decreasing variants, while positive selection to introduce new affinity-increasing amino acid variants is associated with higher ω_CDR_ values. When comparing TNs to NNs we found an overall trend towards fewer amino acid changes in the CDRs of TNs (Maximum likelihood estimate (MLE) of ω_CDR_ in Fig. 3C) indicating stronger purifying selection. In contrast, the average lineage tree lengths (the total expected substitutions per codon site within an individual lineage phylogeny) was significantly higher in TNs compared to NNs (*p* < 0.05 Wilcoxon rank sum test) and correlated negatively with ω (Fig. 3D). The largest lineage trees constructed in TNs and NNs are visualized in Figure 3E and Figure 3F respectively. These results are therefore consistent with prior observations that B cell lineages shift toward negative selection over time as a general feature of affinity maturation (29) and suggest that the dominant clones in the TNs have reached a peak of maximum affinity maturation where further random amino acid changes would rather be detrimental and decrease affinity (30).

## Discussion

Overall, when comparing all paired IGH and IGL sequences, our repertoire analysis did not reveal substantial differences between those individuals who developed antibody neutralization breadth compared to those who did not. In contrast, certain genes that are distinctive for known bNAbs, like VH1-2 or VH1-46 for the CD4bs antibodies (31, 32), HV4-59 for the V3-glycan dependent antibodies PGT121 and 10-1074 (9), or HV3-33 for the V1/2 glycan specific antibodies PG9 and PG16 (9, 33), were present at similar frequencies in TNs or NNs, suggesting, that the prevalence of specific Ig gene segments alone is not predictive of neutralization breadth. Scheepers et al. who analyzed the IGHV repertoires of bulk peripheral blood B-cells for 28 HIV-1 clade C infected South African women with different ranges of serum neutralization breadth (34) observed a wide range in the number of IGHV alleles in each individual, including alleles used by known broadly neutralizing antibodies, but also did not find significant differences in germline IGHV repertoires between individuals with or without broadly neutralizing antibodies. Our analysis not only confirms these findings on an antigen-specific BCR repertoire level but also in the setting of persistently low HIV antigenemia, reducing potential bias due to difference in viral exposure and diversity.

While the composition of the BCR repertoire alone failed to identify TNs, both groups were clearly distinguishable by the significantly higher rates of mutations in the VH and VL genes in the individuals with neutralizing antibody breadth. Our data in HIV-1 specific B-cells confirms findings recently reported by Roskin et al. in a bulk B-cell repertoire analysis, demonstrating that individuals with neutralizing antibody breadth have hundreds to thousands of antibody lineages with long CDRH3s and very high SHM frequencies (27). Moreover, the mutation frequency in our data correlated directly with the neutralization breadth suggesting a skewing towards poly- or cross-reactive antibodies that are able to bind mutated forms of the original epitope. Initial interclonal and then intraclonal competition between B-cells during affinity maturation is well described (35, 36), resulting in diverse repertoires. Indeed, we found that with increasing numbers of mutations in VH, fewer clones with less diverse VH-VJ combinations dominated, suggesting a process that allows these clones to mature and accumulate mutations. Since GCs select for affinity and not for neutralizing breadth and since evolving viruses, such as HIV-1, will continously elicit novel immune responses from naive B cells, strong effective selection normally occurs, so that antibodies specific to the contemporaneous virus have a sufficient advantage to outcompete any less-specific potential broad-neutralizers before they take hold. Indeed, by focusing on the kinship relations among Ig gene segment recombinations within individual clones, we found evidence of natural selection in the NNs, with younger and less diverse clones continuously evolving, constantly exploring novel evolutionary space. Conversely, in TNs, this continuous evolution does not seem to occur, but rather specific evolutionary paths are pursued. One potential mechanism behind this observation is that neutralizers may generate massive bursts of clones that compete aggressively, and prevent the evolution of lower affinity variants. Alterantively, TN clonal diversification may selectively prevent high-affinity antibodies specific for the contemporaneous virus from excluding the broad-neutralizers as they develop. The effects of immunodominance where B-cell specific to an epitopic site dominate B-cells that target other sites, as has been described in influenza infection (37), might not occur to the same degree in TNs as it might do in NNs. Indeed, a generalized deficiency of normal selection against B cells expressing IgG with long CDRH3 regions and high SHM frequencies has been associated with the development of HIV neutralizing breadth (27).

While the selection process might be suppressed in the TNs early on, to allow extended SHM with ongoing clonal evolution and increasing affinity, this process seems to revert during the later stages of antibody evolution when the benefit of additional amino acid changes decreases, and purifying selection becomes dominant and removes most non-synonymous mutations. Indeed, Yaari et al. demonstrated that early mutations in the trunk of B-cell clonal lineages from healthy subjects were consistent with positive selection pressure than more recent mutations in the canopy (38) and a negative relationship between both signs of increased negative selection (low ωCDR values) and mean repertoire tree length have been reported across subjects of different age and sex (29). Interestingly, Wu and Sheng et al observed that the evolutionary rates of HIV-1 bNAb antibody lineages derived from different donors consistently decreased over time (39, 40), also suggesting that SHM occurs at a faster rate during the early phase of lineage development while later on, a decrease in mutation rates may be a mechanism to protect immune memory.

Several potential mechanisms for differences in BCR evolution and selection can be envisioned in TNs versus NNs. B cells with the highest-affinity receptors tend to acquire the most antigen from the follicular dendritic cell (FDC) network and present the highest density of cognate peptides to CD4+ T-follicular helper (Tfh) cells, which respond with survival signals to the B cell. While some studies have suggested that the quality and quantity of germinal center Env-specific Tfh cells are associated with the expansion of Env-specific B cells and broader antibody neutralization activity (41, 42), others have failed to find a correlation (27). Interestingly, Cirelli et al recently demonstrated that slow delivery of antigen resulted in enhanced autologous tier 2 nAb development in non-human primates (NHP) by inducing higher frequencies of total and Env-specific GC-Tfh cells accompanied by larger and more diverse Env-specific B cell lineages. In this model, evolving B cells with BCRs able to bind to a large array of ENV variants, are likely able to capture more antigen, present more effectively to Tfh and thus gain higher survival signals. Conversely, B cells able to bind tightly, but to only a narrow range of ENV variants will only capture a fraction of arrayed antigens, and receive less help. Thus, high affinity, narrow specificities may gain less support compared to lower-affinity, broad-binders (43). Thus, while antigenemia is lower in controllers, the low level titration of antigen into the system, may create a higher bar of selection for evolving B cell clones. Along with this high degree of competition, B cell lineages in HIV controllers able to bind many variants may capture larger amounts of antigen and therefore gain enhanced signals from Tfh (44). Additionally, HIV controllers with low level viremia also exhibit unique inflammatory profiles, and sustained GC activity, collectively also potentially providing an optimal environment for B cell development.

In summary, development of B-cell lineages with superior neutralization breadth in HIV controllers is linked to selected clonal evolution, therefore allowing B-cells to accumulate mutations and to diversify without the constant risk of clonal interference. Immunization strategies that facilitate this process by providing continuous, diverse and subtherapeutically dosed antigen, in the setting of target inflammatory signals, could potentially accelerate the development of broad nAb responses in vaccinees.

## Acknowledgments

We would like to thank the participants of the Ragon HIV controller studies from whom these samples were obtained, and the staff involved in all the sample collection and processing. We thank Dr Daniel Lingwood for the careful review of the manuscript and his critical input. This work was funded by Bill and Melinda Gates Foundation Collaboration for AIDS Vaccine Discovery (CAVD) grant #OPP1146996.

## Author Contributions

MSS generated the antibody neutralization data. AS, DK, NN, YCT sorted antigen-specific B-cells. DK, NN, YCT and GC performed single cell BCR sequencing, data quality control and sequence analysis. DC and GL performed the BCR analysis and GA and BJ supervised the analysis. DC, GA and BJ wrote and all authors reviewed the manuscript.

## Declaration of Interests

DK, GC, NN are employess of Atreca Inc. YCT was an employee of Atreca Inc. during the generation of the data and is now an employee of Esco Ventures X. GA is a founder of Seromyx Systems. The other authors declare no conflict of interests.

## Supplemental Material

**Supplemental Table 1:**
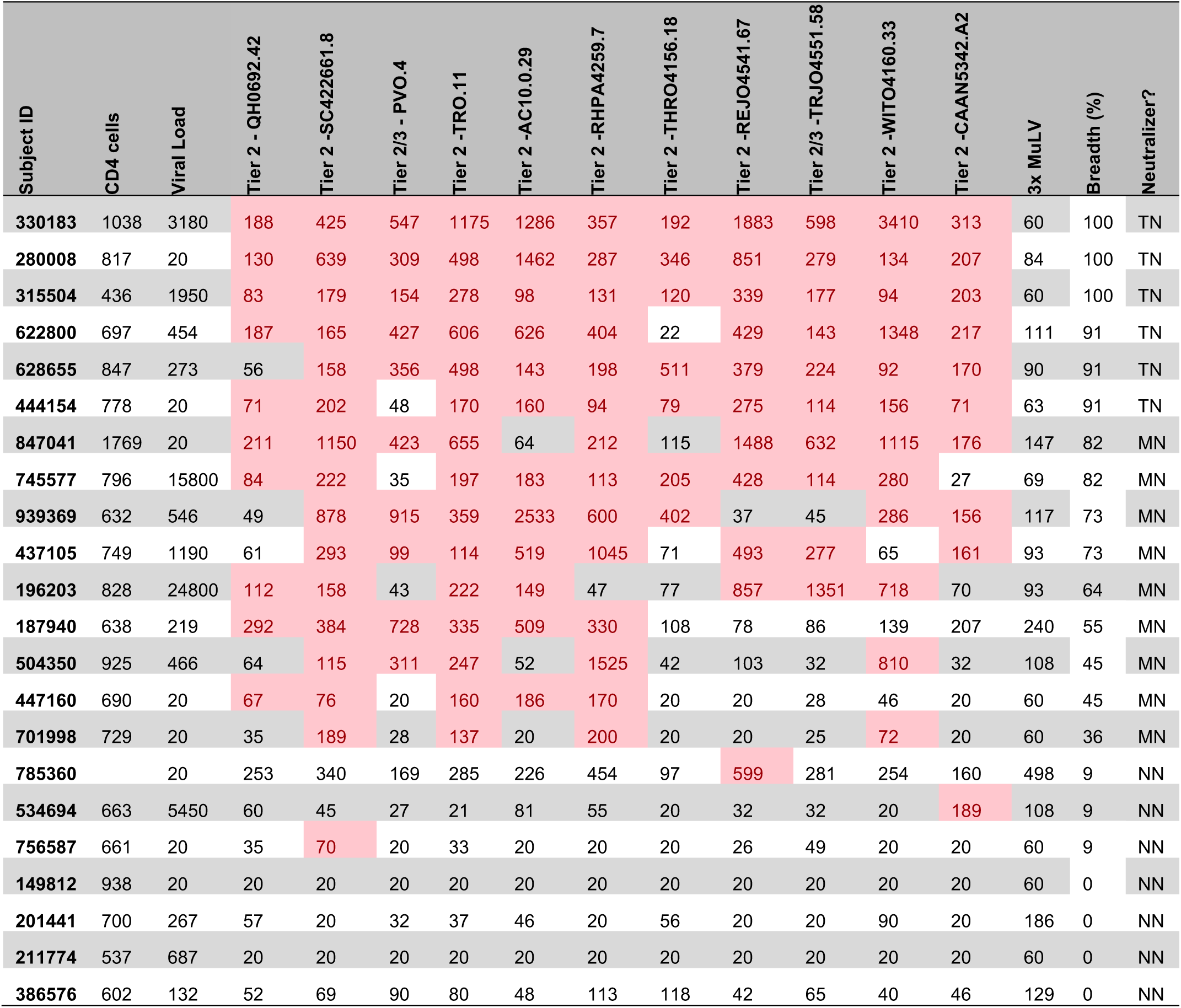
Study participant characteristics. CD4 T-cell counts (cells/ul), viral loads (HIV RNA copies/mL) and neutralization titers [50% inhibitory dose (ID50), 1/x] from each individual against against a panel of eleven tier 2 and 3 Env-pseudoviruses and murine leukemia virus– pseudotyped virion (MuLV) control. The right column (Breadth %) shows the % of viruses that are neutralized above background (3x MuLV control). Six individuals Neutralization of more than 90% of the panel classified as Top-Neutralizer (TN) while neutralization of less than 10% of the panel viruses classified as Non-Neutralizer (NN).

**Supplemental Figure 1:**
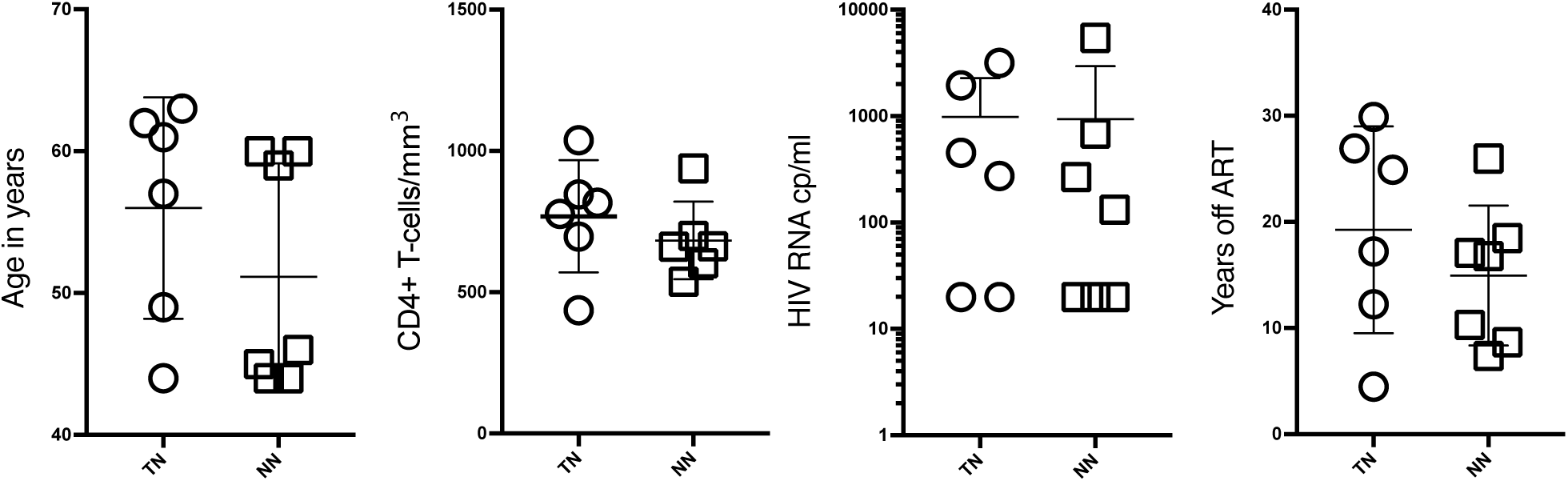
Study participant characteristics at time of sampling.

**Supplemental Figure 2:**
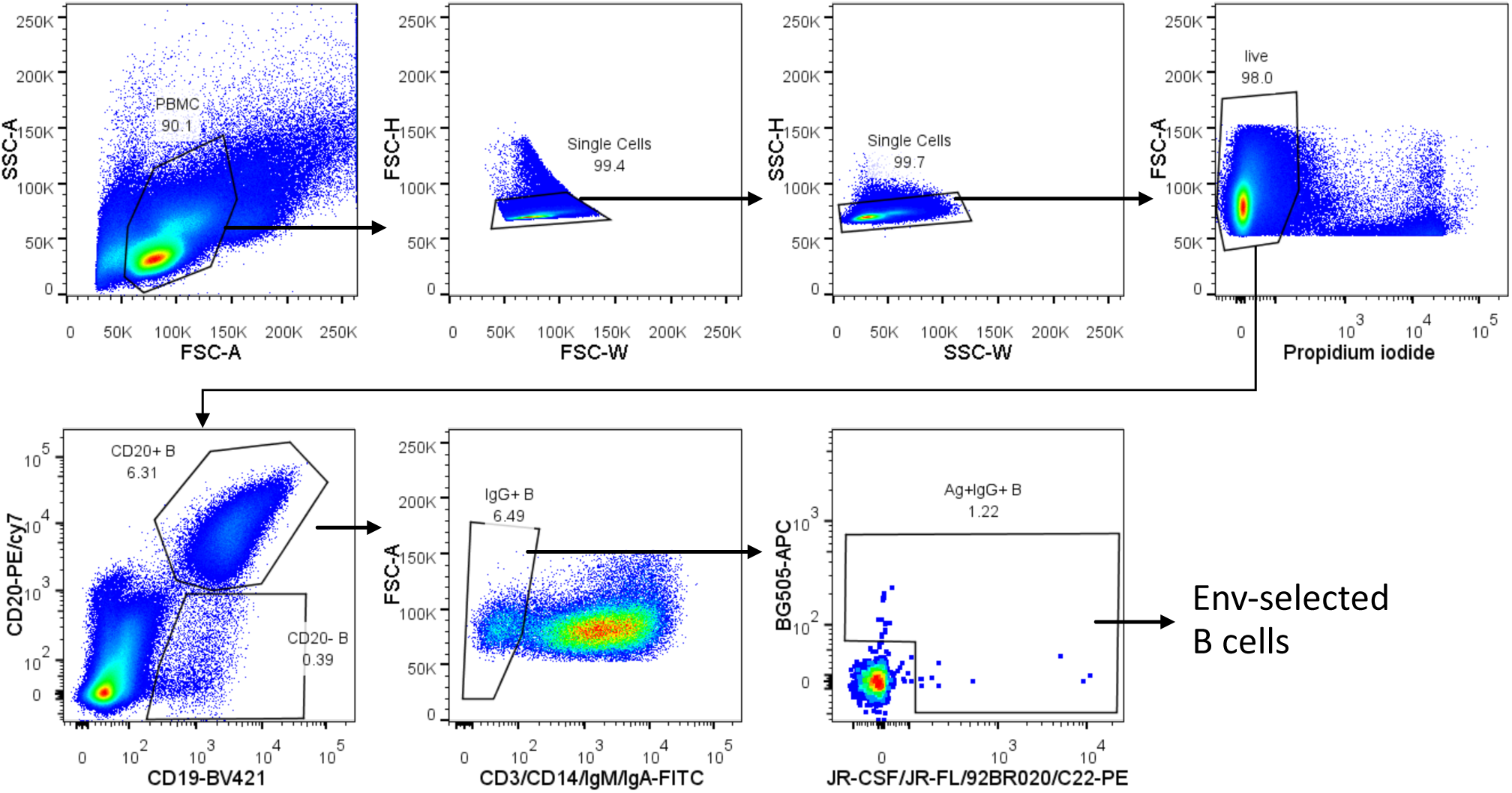
Gating strategy to identify and sort single Env-specific memory B cells.

## Methods

### Human subjects

A total of 22 HIV-infected individuals were recruited for this study. All participants underwent leukapheresis and routine blood draws as per study protocol. All subjects signed informed consent, and the study was approved by the MGH/Partners Institutional Review Board (Protocol #: 2003P001894).

### Neutralization Assay

HIV-1 neutralization breadth was assessed using the Tzm-bl cell–based pseudovirus neutralization assay, as described (45), against a standard panel of Env-pseudoviruses derived from 9 Clade B Tier 2 viruses: AC10.0.29, RHPA4259.7, THRO4156.18, REJO4541.67, WITO4160.33, TRO.11, SC422661.8, QH0692.42, CAAN5342.A2, and 2 Tier 3 viruses: PVO.4 and TRJO4551.58 (12). Murine leukemia virus (MuLV) was included in all assays as a negative control. Neutralization titers [50% inhibitory dose (ID50)] were defined as the reciprocal of the plasma sample dilution that caused a 50% reduction in RLUs compared to virus control wells after subtraction of background RLUs. Neutralization breadth was determined as the proportion of pseudoviruses with an ID50 score 3-fold above background titers observed against MuLV negative control virus (3x ID50 of MuLV).

### Single-cell flow cytometry sorting

Isolated cells were stained with fluorochrome-antibody conjugates and reagents to identify antigen-specific memory B cells. The panel consisted of propidium iodide (Life Technologies); CD3 (FITC, clone UCHT1), CD14 (FITC, clone HCD14), IgM (FITC, clone MHM-88), CD20 (PE-cy7, clone 2H7), CD19 (BV421, clone HIB19) (All Biolegend); IgA (FITC, clone IS11-8E10) (Miltenyi Biotec). Preformed conjugates for antigen-specific B cell sorting were made as described (46) using streptavidin conjugated to PE or AlexaFluor 647 (Life Technologies). B cell probes were made using clade B JR-CSF gp120, JR-FL gp140, 92BR020 gp120, clade A BG505 SOSIP, and clade C IAVI C22 gp120 tags (Duke Human Vaccine Institute protein production fascility). IgG+ B cells were defined as CD3/14−, CD19+, CD20+, and IgA/IgM-; antigen-specific B cells positive for probes in either PE or APC color were sorted into 96-well U-bottom plates containing 200 µl of B cell culture medium (IMDM supplemented with FBS, Normocin, hIL-2, hIL-21, hCD40L His tag, and anti-His antibody). After 4-day culture, the B cells were sorted into microtiter plates at one cell per well. Sorted plates were frozen immediately and maintained at −80C before RT/PCR.

### B-cell receptor sequencing

Natively-paired variable region sequences from individual cells were generated by reverse transcription, cDNA barcoding, amplification and sequencing as described previously (47, 48). cDNA sequences were determined by 454 Titanium sequencing. A minimum of 10 reads for each chain (heavy and light) was required, and a contig was kept only if it included at least 90% of the reads for that chain from that well. V(D)J assignment and mutation identification was performed using an implementation of SoDA (49).

### Analysis

Statistical analyses and visualizations were performed in R version 4.0.0 (2020-04-24). Subclass distribution, gene family and gene usage analysis: To exclude potential bias caused by the number of input cells or sequences, we divided the number of occurances of each repertoire signature by the total number of all sequences for each individual. IGHV-J recombination profiles: to avoid skewing of mean frequencies when averaged across all individuals by individuals with high or low number of available sequences (in particular for sequences with more than 60 heavy chain mutations) normalized frequencies were multiplied by a factor (log2 of total sequences). To calculate CDRH3 length distribution, all TN and all NN sequences were combined.

Mean percantages of gene usage between TN and NN were compared using two-way ANOVA and p-values were corrected for multiple comparison using Sidak’s post-hoc test. Mean number of mutations between TN and NN were compared using unpaired two-samples t-test. All correlations were performed using Spearman’s correlation. Per each TN subject, the enrichment of V-J combinations in MBCs with high mutation rates were tested using one-sided Fisher exact test. Visualizations were performed using R package ggplot (version 3.3.0). Circos plots were generated using R package circlize (version 0.4.9) *chordDiagram* function. The layout for clonal network visualization was arranged using R package packcircles (version 0.3.3).

Clones were defined as cluster of sequences that meet the following criteria (1) come from the same individual; (2) share the same IGHV and IGHJ gene segment annotations (3) have equal CDRH3 length; and (4) CDRH3 aa sequences match with 90% or more similarity. CDRH3 sequence homology was calculated using R package Biostrings (version 2.56.0) *pairwiseAlignment* function. The clonality measure was estimated using Gini index and calculated using R package bcRep (version 1.3.6) *clones.giniIndex* function.

Lineage trees were contructed using IgPhyML (version 1.1.0)(50). The HLP19 model was used to estimate maximum likelihood tree topologies. Clonal selection was estimated using ω: Also called dN/dS, or the ratio of nonsynonymous (amino acid replacement) and synonymous (silent) mutation rates. A value of ω≈1 indicates totally neutral amino acid evolution, ω<1 negative selection and ω>1 diversifying selection. The topology analyses were carried out using Alakazam (version 1.0.0) (51) and the lineage trees were visualized using R ggtree (version 2.2.1).

